# Symmetry breaking and fate divergence during lateral inhibition in *Drosophila*

**DOI:** 10.1101/2024.03.11.583933

**Authors:** Minh-Son Phan, Jang-mi Kim, Cara Picciotto, Lydie Couturier, Nisha Veits, Khallil Mazouni, François Schweisguth

## Abstract

Lateral inhibition by Notch mediates the adoption of alternative cell fates amongst groups of initially equipotent cells, leading to the formation of regular patterns of cell fates in many tissues across species. Genetic and molecular studies have established a model whereby an intercellular negative feedback loop serves to amplify small stochastic differences in Notch activity, thereby generating ordered salt-and-pepper patterns. In *Drosophila*, lateral inhibition selects Sensory Organ Precursor cells (SOPs) from clusters of proneural cells that are competent to become neural through the expression of proneural transcription factors. When and how symmetry breaking occurs during lateral inhibition remains, however, to be addressed. Here, we have used the pupal abdomen as an experimental model to study the dynamics of lateral inhibition in *Drosophila*. Using quantitative live imaging, we monitored the accumulation of the transcription factor Scute (Sc), used as a surrogate for proneural competence and adoption of the SOP fate. We found that fate symmetry breaking occurred at low Sc levels and that fate divergence was not preceded by a prolonged phase of low or intermediate level of Sc accumulation. The relative size of the apical area did not appear to bias this fate choice. Unexpectedly, we observed at low frequency (10%) pairs of cells that are in direct contact at the time of SB and that adopt the SOP fate. These lateral inhibition defects were corrected via cellular rearrangements. Analysis of Sc dynamics in wild-type and genetically mosaic pupae further revealed that cell-to-cell variations in Sc levels promoted fate divergence, thereby providing experimental support for the intercellular negative feedback loop model.

## Introduction

During development, stereotyped patterns of cell fates can be produced via self-organization guided by positional cues (Green and Sharpe, 2015)(Schweisguth and Corson, 2019). Lateral inhibition is a conserved self-organized patterning process whereby equipotent cells inhibit each other from adopting the same fate via inhibitory cell-cell interactions mediated by Notch receptor signaling (Simpson, 1990)(Bray, 2006)(Kopan and Ilagan, 2009)(Henrique and Schweisguth, 2019). In the absence of guiding cues, lateral inhibition produces ‘salt-and-pepper’ patterns of cell fates, but more elaborate patterns can be generated when a prepattern of Notch activation serves as initial condition for lateral inhibition (Corson et al., 2017). Classic studies in *Drosophila* have indicated that stochastic fate decisions rely on the amplification of random fluctuations in Notch and/or proneural activity via an intercellular negative feedback loop (Fig 1A) (Heitzler and Simpson, 1991)(Simpson, 1997)(Greenwald and Rubin, 1992)(Heitzler et al., 1996). In addition, mathematical modeling showed that an intercellular feedback loop can produce salt-and-pepper patterns when the negative feedback is strong enough to amplify fluctuating differences between adjacent cells (Collier et al., 1996). In *Drosophila* sensory organ formation, this feedback loop has four key elements (Fig 1A’): the receptor Notch, its ligand Delta (Heitzler and Simpson, 1991), the E(spl)-HLH factors encoded by the direct transcriptional targets of Notch (Bailey and Posakony, 1995)(Lecourtois and Schweisguth, 1995)(Delidakis et al., 2014)(Heitzler et al., 1996)(Castro et al., 2005), and the transcription factors Achaete (Ac) and Scute (Sc) (Skeath and Carroll, 1991)(Cubas et al., 1991)(Campuzano and Modolell, 1992). Ac and Sc specify the fate of the Sensory Organ Precursor cells (SOPs) and are antagonized by Notch/E(spl)-HLH. They also positively regulate the signaling activity of Delta, possibly via an indirect mechanism (Seugnet et al., 1997)(Pitsouli and Delidakis, 2005)(Chanet et al., 2009). While the regulatory logic and general relevance of this feedback loop are well established (Sánchez-Iranzo et al., 2022)(Simpson, 1997)(Castro et al., 2005)(Hellström et al., 2007)(Apelqvist et al., 1999), the spatial and temporal dynamics of both Notch signaling and proneural activity during lateral inhibition are still largely unexplored. Indeed, the time-course expression of Ac and Sc and the dynamics of fate acquisition has mostly been studied in fixed samples in *Drosophila* (Skeath and Carroll, 1991)(Huang et al., 1991)(Corson et al., 2017; Couturier et al., 2019)(Cubas et al., 1991)(Usui and Kimura, 1993)(Couturier et al., 2019)(Troost et al., 2015). These studies indicated that groups of cells first accumulated intermediate levels of proneural factors before one presumptive SOP was detected on the basis of increased levels of Ac and Sc, and that SOP selection was accompanied by the down- regulation of Ac and Sc in the surrounding cells that were inhibited by Notch. However, when, and how symmetry breaking (SB) occurs during lateral inhibition remained undefined. Here, we studied in real time the dynamics of cell fate decision during lateral inhibition by live imaging a functional GFP-tagged version of the proneural factor Sc (Corson et al., 2017). We focused our analysis on the selection of SOPs in the developing abdomen because high-resolution imaging can be easily performed around the time of SOP specification (Ninov et al., 2009)(Davis et al., 2022). Quantitative image analysis allowed us to identify SB and measure the rate of fate divergence during lateral inhibition. This showed that fate selection occurred early and was not preceded by a detectable phase of mutual inhibition, and that cell-to-cell variations in Sc levels at SB promoted fate divergence.

**Figure 1:**
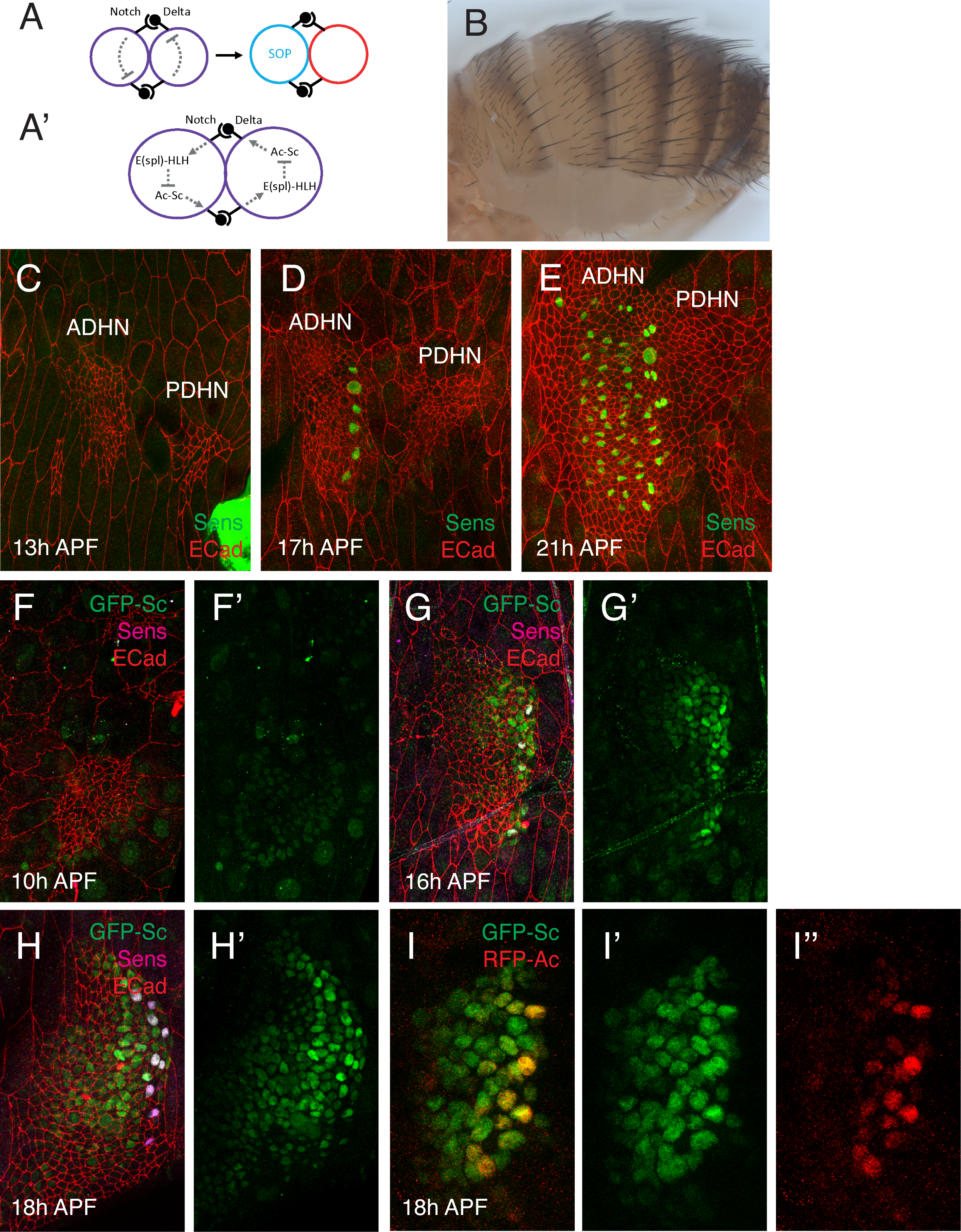
Cell fate patterning. A,A’) Lateral inhibition relies on an intercellular negative feedback loop whereby Notch activation in a given cell inhibits Delta signaling activity in the same cell (A); four core elements form this feedback loop (A’). B) Lateral view of an adult fly showing the stereotyped distribution of sensory bristles in the abdomen, with a posterior row of large bristle and a disordered array of smaller bristle at the center. In this and all other images, dorsal is up and anterior is left. C-E) Time-course of SOP emergence in the pupal abdomen. SOPS were marked with Senseless (Sens, green). The ADHN and posterior dorsal histoblast nest (PDHN) are seen as clusters of small diploid cells (E-Cad, red). SOPs were detected first along the posterior margin of the ADHN (D), then in the central region (E). The ADHN and PDHN fused after the emergence of these posterior SOPs (E). F-H’) Time-course of GFP-Sc expression (GFP-Sc, green) in the ADHN (E-Cad, red). SOPs were marked with Sens (magenta). I-I’’) RFP-Ac (red) was detected in emerging SOPs and in a small number of neighboring cells. In contrast GFP-Sc (green) was broadly expressed in the ADHN.

## Results

### Time-course analysis of SOP emergence in the pupal abdomen

Adult sensory bristles are distributed in two distinct patterns at the surface of the dorsal abdomen (Fig 1B): large sensory bristles appear at regular interval along the compartment boundary to form a posterior row whereas smaller bristles are distributed in a disordered array more anteriorly. Confirming earlier reports (Shirras and Couso, 1996)(Davis et al., 2022), these two patterns were detected at the time of SOP emergence in the pupal abdomen: SOPs producing large sensory bristles appeared first along the posterior edge of the Anterior Dorsal Histoblast Nest (ADHN), while more anterior SOPs emerged later in the central region of the ADHN (Fig 1C-E). Since both types of SOPs depend on the proneural genes *achaete* (*ac*) and *scute* (*sc*) for their determination (Fig S1), we examined the expression pattern of these factors using tagged version of Sc and Ac (Corson et al., 2017). GFP-Sc was not expressed prior to 14h after puparium formation (APF; 10h APF in Fig 1F,F’) and was first detected in a broad stripe of cells located along the posterior margin of the ADHN (16h APF in Fig 1G,G’). The pattern of GFP-Sc expression then progressively extended in the central region of the ADHN (18h APF in Fig 1H,H’). In contrast with GFP-Sc that was broadly expressed in the ADHN, RFP-Ac expression was restricted to emerging SOPs and surrounding cells (Fig 1I-I’’), indicating that that Ac is activated later than Sc and in fewer proneural cluster cells. Since Sc, but not Ac, was expressed broadly in the proneural field prior to SOP selection, we focused below on the dynamics of Sc accumulation during lateral inhibition.

### Live imaging of Scute

To study the dynamics of SOP emergence, we performed live imaging using GFP-Sc. Staged pupae were imaged from ∼14h APF, starting prior to the onset of GFP-Sc, to ∼22-24h APF, ending after all SOPs have been specified (Fig 2A-D’, movie 1). In these movies, SOPs were identified as isolated cells with high GFP-Sc intensity at ∼20-22h APF (Fig 2D’’). The intensity of the nuclear GFP-Sc signal was measured over time in (x,y,z,t,c) movies (n=3). A nuclear RFP was used to segment and track individual nuclei. Image processing involved denoising and 3D segmentation of nuclei followed by 3D object filtering to eliminate bright dots corresponding to fragmented apoptotic nuclei as well as large fluorescent structures (see Methods). We first examined the dynamics of Sc expression at the tissue-scale by projecting along the A-P axis the GFP-Sc intensity values from all cells to produce kymographs (Fig 2E). This analysis confirmed that GFP-Sc appeared first along the posterior margin of the ADHN, then in the central domain of the ADHN. Our observations are consistent with earlier findings suggesting that posterior cues might regulate early proneural gene expression in the ADHN (Shirras and Couso, 1996)(Kopp et al., 1997)(Bischoff et al., 2013). To study fate dynamics, SOPs from the central region of the ADHN were then tracked in 3D, both forward (until they divide asymmetrically to produce a pIIa-pIIb cell pair) and backward (up to their birth from a dividing histoblast progenitor cell). A total number of 130 SOPs were tracked. Following manual curation of all tracks, normalized GFP-Sc levels were measured in tracked SOPs. The temporal expression profile of GFP-Sc in tracked SOPs relative to all other untracked cells indicated that SOPs emerge within a ∼2h time window centered around ∼17h APF in the ADHN (Fig 2F).

**Figure 2:**
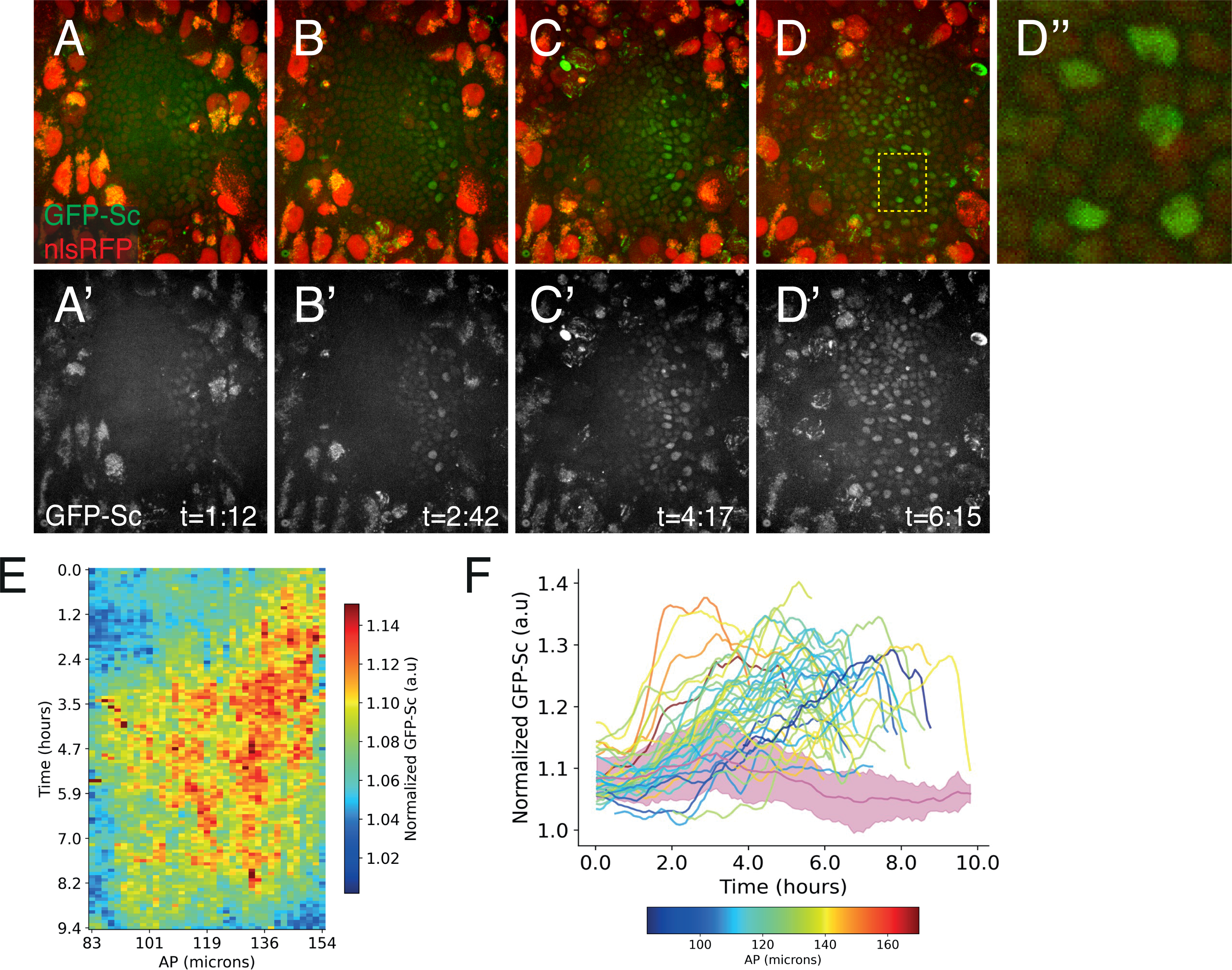
Scute dynamics. A-D’’) Live imaging of GFP-Sc (green; nlsRFP, red) in the pupal abdomen (see movie 1). SOPs were identified as isolated cells with high levels of GFP-Sc (D’’, high magnification view of the region boxed in D’). Note the strong autofluorescence signal in the larval epidermis. E) Kymograph showing the temporal pattern of GFP-Sc accumulation (color-coded intensity) plotted along the A-P axis. GFP-Sc was first expressed in posterior ADHN cells, then in the central region. F) Temporal profile of GFP-Sc expression in tracked SOPs (color-coded based on their position along the A-P axis). SOPs emerged first in the posterior row at ∼15-16h APF (∼1-2h after the start of imaging), then in the central domain at ∼16-18h APF. The temporal profile of GFP-Sc measured in non- SOP cells is indicated in pink. Both the mean and standard deviation (std) are shown.

### Fate symmetry breaking at low Sc levels

We next studied the dynamics of cell fate acquisition by comparing the dynamics of Sc accumulation in each presumptive SOP relative to its immediate neighbors, defined here as the six closest nuclei, in a three-dimensional space, following manual correction of segmentation errors (Fig 3A,A’; note that in the absence of cortical markers, we assumed but could not demonstrate that these neighboring nuclei marked cells that were in direct contact with the SOP). This number of neighbors was chosen because we found that SOPs had 5.8 +/- 1.1 apical neighbors (n=35) between 17 and 18h APF in GFP-tagged E-Cadherin (CadGFP) movies (see methods and Fig 5 below). Since these nuclei were not tracked, different nuclei could participate over time to the cluster of cells associated with a given SOP. A representative example of the temporal profiles of GFP-Sc levels in a single SOP and in its six neighbors is shown (Fig 3B). In this example, the presumptive SOP and its surrounding histoblasts showed a weak and slowly increasing GFP-Sc signal from t=1.5h onwards until the presumptive SOP showed a rapid increase in GFP-Sc accumulation at ∼2.8h (Fig 3B; this experimental time corresponds to ∼17h APF). As GFP-Sc levels increased sharply in the SOP, GFP-Sc levels remained relatively constant in non-selected histoblasts for more than one hour before decreasing. This temporal profile suggested that cells underwent binary fate decisions around the time when GFP-Sc started to increase in the presumptive SOP. To study more quantitatively fate divergence, we introduced a fate difference index (FDI) that compares the relative level of Sc in the presumptive SOP and in its closest neighbors. This FDI was defined as the ratio of the difference over the sum of the GFP-Sc intensities measured in the SOP nucleus and in the six closest nuclei. This allowed us to detect when the SOP becomes different from its neighbors (Fig 3C). To study the onset of fate divergence, corresponding to fate symmetry breaking (SB), we defined a time point when the FDI value reached and remained above a given threshold. To set the value of this threshold value, we measured FDI values over time from groups of randomly sampled non-SOP cells. This analysis indicated that background FDI values associated with cell-to-cell differences in GFP-Sc levels in non-SOP cells were below 0.005 (Fig S2). We therefore chose this threshold value to define SB. This minimal FDI value was found to be superior to the fluctuations observed before the sharp increase in GFP-Sc levels, and we found that FDI values consistently increased after SB for almost all tracks (Fig 3C). We also used the FDI to quantitatively evaluate fate divergence, using the Rate of Change (ROC) of the FDI as a proxy for the rate of fate divergence, or speed of fate decision.

**Figure 3:**
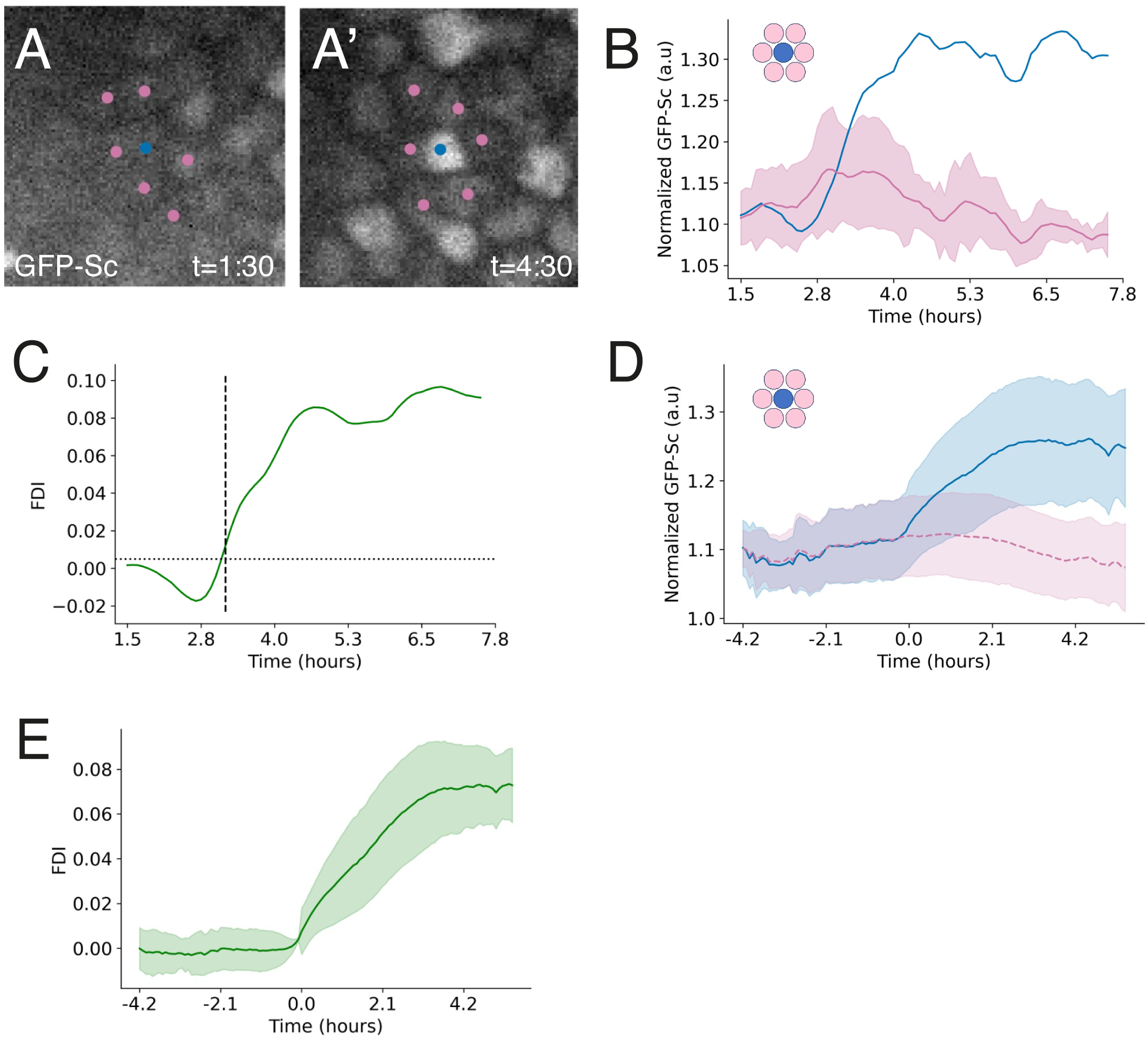
Fate symmetry breaking. A-B) GFP-Sc accumulation in a single tracked SOP (blue dot in A,A’; blue curve in B) and in its six closest nuclei (in 3D, not tracked; pink dots in A,A’; pink curve in B showing the mean and std was plotted over time (B). GFP-Sc accumulation before and after SOP emergence is also shown in 2D projected views of the cluster (A,A’). C) Temporal profile of the Fate Difference Index (FDI) for the cluster shown in panels A-B. Positive values indicate that the level of GFP-Sc in the future SOP is on average higher than in its close neighbors. A Symmetry Breaking (SB) point was defined as the time when FDI is stably above a threshold set to 0.005 (horizontal dashed line). The SB point is indicated by a vertical dashed line. D) Normalized GFP-Sc intensities in SOPs (blue) and neighboring cells (pink) for all clusters (n=130) that were registered in time using SB as a reference. SB was observed early at low level of GFP-Sc. E) Temporal profile of FDI values for all registered clusters. A clear linear increase of fate divergence was observed from the time of SB onwards.

To go beyond this example and evaluate more systematically cell fate dynamics, we used SB as a reference time point to re-align all tracks (n=130). This analysis revealed that the onset of fate divergence was observed at low GFP-Sc levels, before presumptive SOPs accumulate high levels of GFP-Sc (Fig 3D). It further showed that the FDI increased at a relatively constant rate after SB, reaching a plateau 3-4h after SB (Fig 3E). However, we did not observe a phase during which many proneural cluster cells would express intermediate level of Sc prior to fate selection, which might correspond to the proposed phase of mutual inhibition (Cubas et al., 1991)(Muskavitch, 1994). Also, we did not observe a rapid decrease of proneural expression in non-selected cells after SB. Instead, the level of GFP-Sc remained relatively constant in non-selected cells for ∼2h after SB. Thus, our analysis of the temporal dynamics of cell fate acquisition in the pupal abdomen showed that fate symmetry breaking takes place at low Sc levels, soon after the onset of proneural gene expression.

### Cell-to-cell variations in Sc levels promote fate divergence

We next studied the heterogeneity of Sc, or cell-to-cell variations in Sc levels, defined here as the coefficient of variation (standard deviation over the mean) of GFP-Sc intensities measured in the presumptive SOP and its six neighbors. Analysis of heterogeneity over time showed that it began increasing ∼1h before SB and remained high in non-selected histoblasts for about 2h before decreasing down to initial levels (Fig 4A). Interestingly, the speed of SOP emergence, as determined using the ROC of FDI, positively correlated with heterogeneity of Sc at SB (Fig 4B). These observations raised the possibility that heterogeneous Sc levels in proneural clusters might promote fate decision. To test this idea, we experimentally increased the heterogeneity of Sc by creating marked clones of cells expressing a RNAi against GFP (Fig 4C-C’’; loss of *sc* activity did not prevent SOP formation, Fig S1). Clone border analysis confirmed that cells with reduced proneural activity were less likely to become selected as SOPs (Heitzler et al., 1996) (Fig S1). We then studied by live imaging the dynamics of GFP-Sc in these genetically mosaic pupae. Following segmentation of nuclei marked by nuclear RFP, we tracked the wild-type SOPs located along the clone boundary (n=165, 12 movies) and identified the six closest SOP neighbors at each time point (Fig 4D,D’). We then plotted the FDI over time (excluding RNAi-expressing cells to measure fate competition between wild-type cells) and determined SB for each cluster. We also determined the mean number of RNAi-expressing cells around the time of SB. As expected, the heterogeneity of Sc measured at SB using all cluster cells, hence including the RNAi-expressing cells, increased with the mean number of RNAi-expressing cells in the cluster (Fig 4E; at least when less than half of the cells, n≤3.5, are expressing the RNAi construct). To test whether increased heterogeneity of Sc in this experimental condition promoted fate divergence, we plotted the ROC of the FDI at SB as a proxy for fate divergence. This analysis showed that increasing the heterogeneity of Sc positively correlated with an increased rate of fate divergence (Fig 4F). Consistent with this, FDI values increased faster in heterogenous clusters (Fig 4G). These results showed that cell-to-cell variations in Sc levels promotes fate divergence during lateral inhibition, hence providing experimental support to the negative feedback loop model that was proposed to amplify small differences and speed up fate divergence (Heitzler and Simpson, 1991)(Heitzler et al., 1996)(Collier et al., 1996). This model also predicted that GFP-Sc dynamics in presumptive SOPs should be locally influenced by Sc dynamics in neighboring cells. Interestingly, late-emerging SOPs are more likely to be surrounded by one or several cells that have low proneural activity due to their inhibition by SOPs that have emerged earlier. We therefore predicted that late-resolved clusters should be more heterogenous, and that late-emerging SOPs should progress more rapidly towards an SOP fate in wild-type pupae. In support of these predictions, we found that both the heterogeneity of Sc and the Rate of Change (ROC) of the FDI positively correlated with the developmental time of SB (Fig 4H,I). Thus, late-emerging SOPs appeared to progress faster towards the SOP fate. Together, our data support the view that initial differences in proneural gene expression become amplified to generate stable cell fates and that cell-to-cell heterogeneity in Sc levels underlies stochastic fate choice in the abdomen.

**Figure 4:**
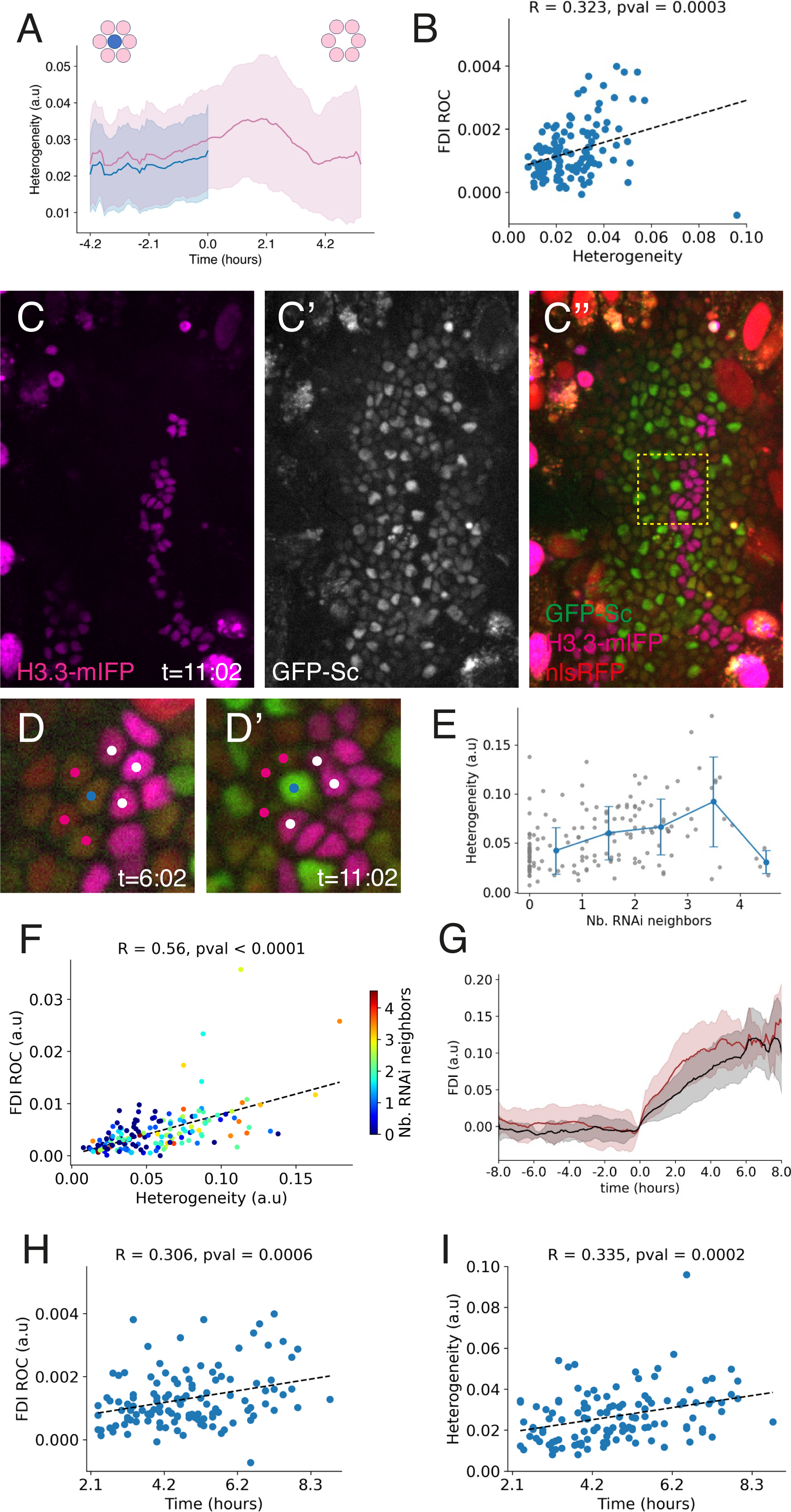
Heterogeneity of Scute and speed of fate decision. A) Quantitative analysis of cell-to-cell heterogeneity in GFP-Sc levels over time. For each cluster, two values were calculated, including the SOP (blue curve; only values prior to SB were plotted because elevated GFP-Sc levels in SOPs led to uninformative increased heterogeneity afterwards) or excluding it (pink curve). B) Correlation between heterogeneity, calculated using all cells of the cluster including the SOP, measured at SB +/- 10min, and the ROC of FDI, measured just after the SB (within 20 min). C-C’’) Snapshot from a movie of a mosaic ADHN (see movie 2). Expression of GFP-Sc (C’, green in C’’; nlsRFP, red) was silenced by a GFP RNAi construct expressed in clones of marked cells (Histone3.3-mIFP, magenta; C). GFP-Sc expression was not detected in the clone. D,D’) Expression of GFP-Sc (green) before and after SOP emergence (D’ corresponds to a high magnification view of the region boxed in C’’). The tracked SOP is indicated with a blue dot; its untracked control and RNAi-expressing neighbors are indicated with magenta and white dots, respectively. E,F) Heterogeneity, calculated around SB +/- 10 min using all cells of the cluster, increased with the number of RNAi-expressing cells reaching a maximum at n=3.5 (E), and positively correlated with the ROC of the FDI at SB (F). G) Temporal profile of the FDI for clusters with low (bottom 50%; in grey) and high (top 50%; in dark red) heterogeneity values. Faster fate divergence was observed in heterogenous clusters. H, I) The onset of SB (developmental time) correlated positively with the ROC of FDI (H), indicating that late emerging SOPs become more rapidly different from their neighbors. It correlated also with the heterogeneity in GFP-Sc levels (I). Thus, early specified SOPs appeared to create local heterogeneity, which in turn promoted fate divergence.

### Relative size of cell-cell contacts does not bias fate decision

We next addressed whether the initial variations of Sc accumulation were random or resulted from some intrinsic bias associated with relative cell-cell contact size. Indeed, earlier modeling suggested that differences in apical cell area may serve as a possible source of bias for Notch-based decisions (Shaya et al., 2017). To test this possibility, we measured apical cell area of abdominal cells in CadGFP pupae that were imaged live. In these movies, SOPs were identified using a nuclear GFP expressed under the control of a SOP-specific enhancer (*pneur-nlsGFP*) (Fig 5A-B’). Following segmentation of apical cell junctions, SOPs were tracked back in time (n=22, 3 movies). Each SOP, together with the untracked cells in direct contact with the SOP, defined a cluster. To test whether apical cell area differ between SOPs and its direct neighbors at the time of fate decision, one would like to determine SB and register in time the different clusters. Unfortunately, SB could not be determined in these movies. However, the time of SB correlated temporally with the division of the SOP in our GFP-Sc nuclear RFP movies and found that SOPs divided ∼4.5h after SB (Fig 5C; no correlation was found between SB and the previous division that generates the future SOP). We therefore registered in time all clusters using as a reference timepoint the SOP division (t=0h in Fig 5D). For each cluster, the apical area of the presumptive SOP was compared to the mean area of its neighboring cells. This analysis revealed no relative difference in apical area around the predicted time of SB (Fig 5D). Thus, apical cell shape did not appear to bias Notch-mediated fate decision in this developmental context.

**Figure 5:**
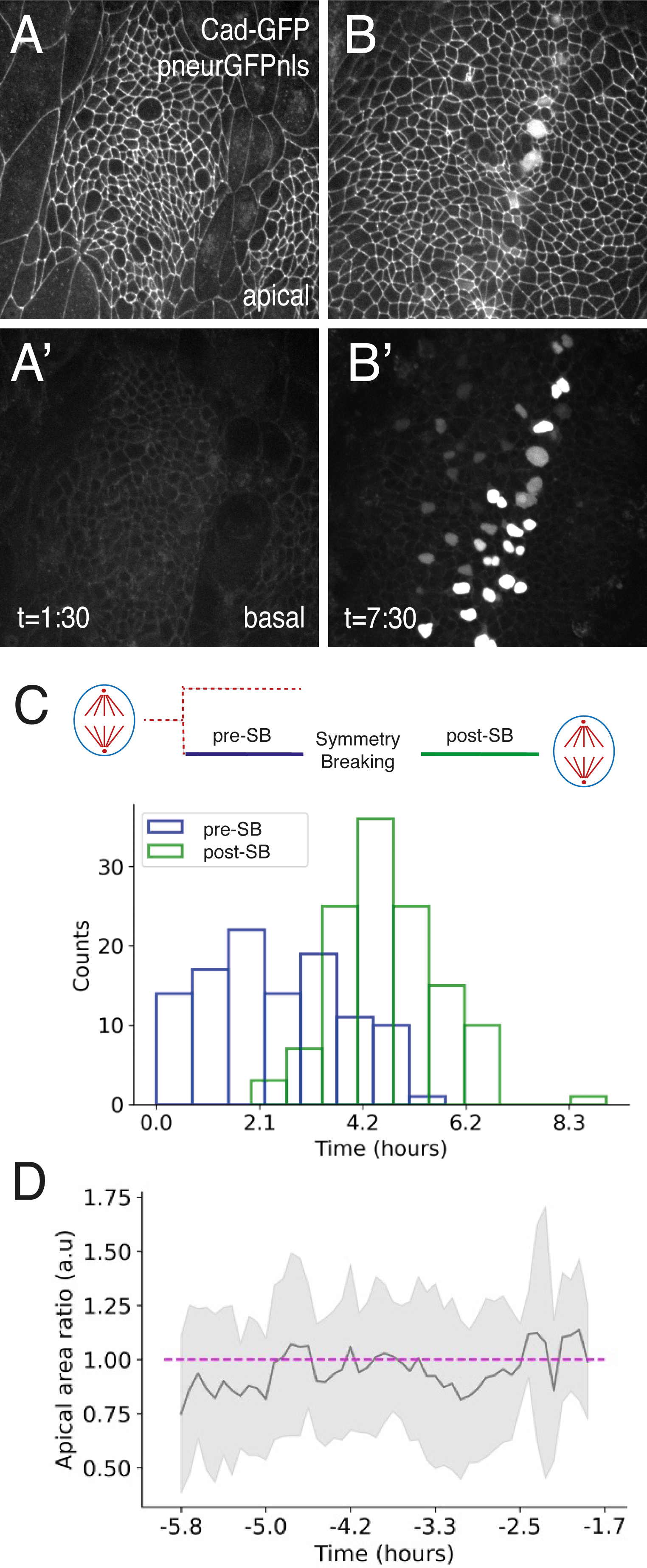
Apical area and cell fate bias. A-B’) Snapshots from the live imaging of a E-Cad-GFP pupae (apical views in A,B) expressing also a nuclear GFP marker in SOPs (basal views in A’,B’; the nlsGFP signal was also detected apically in mitotic cells in B). Two time points are shown (see also movie 3). C) Distribution of the time intervals separating SB from the previous division (pre-SB time) and from the SOP division (post-SB time). These were determined in the GFP-Sc nlsRFP movies shown in Fig 2 (n=125 lineages; in many lineages, only one time interval was scored). This analysis showed that SOPs divided ∼4.5h after SB. No temporal correlation was observed between the previous division and SB. D) Ratiometric analysis of the cell surface area measured over time in each SOP relative to its immediate neighbors (mean value). Each cluster was time registered relative to the SOP division (t=0; SB mapped at -4.5h). No difference in apical area was observed around the time of SB.

### Cell-cell rearrangements correct fate decision defects

While studying Sc dynamics, we observed clusters in which one of the six SOP neighbors had high GFP-Sc levels after SB (n= 16/130) and corresponded to a presumptive SOP located very close to the tracked SOP (Fig 6A-C’). However, in the absence of a cortical marker, it was unclear whether these pairs of SOPs were in direct contact at SB. To address this, we performed live imaging of wild-type GFP-Sc pupae expressing a Cad-mKate marker and identified 20 pairs of SOPs based on high GFP-Sc expression (9 movies; n= 399 SOPs; Fig 6D- E’). Tracking back the apical cell area of these high-GFP scute cells and inferring the time of SB using the SOP division as a time reference (see Fig 5C), we found that most pairs of SOPs (n=18/20) were in direct contact around SB, i.e. before they could be identified as SOPs. Additionally, we observed that these paired SOPs separated from each other at 1h +/-1.7h (n=13/18) after SB, prior to their division. Thus, lateral inhibition in the pupal abdomen failed to single out SOPs in ∼10% of the clusters, and subsequent pattern refinement involved cellular rearrangements following fate specification.

**Figure 6:**
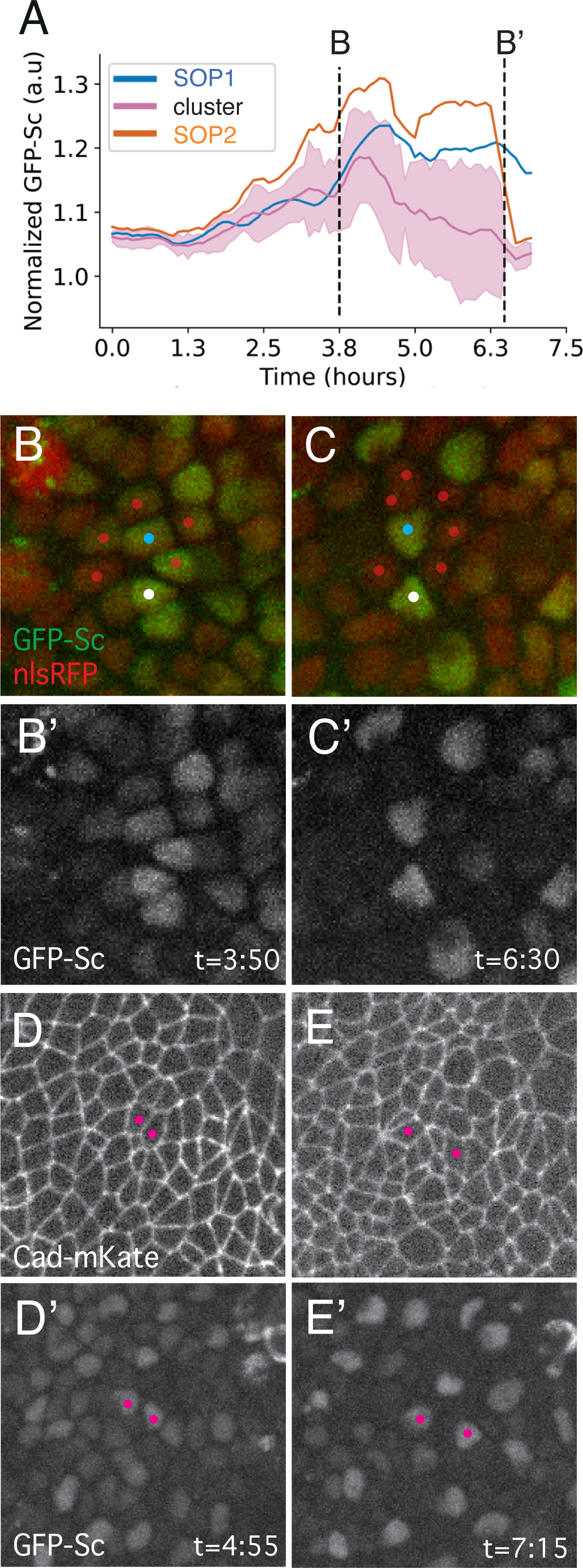
Post-SB patterning through cellular rearrangements. A-C’) Temporal profile of normalized GFP intensities measured over time in cells of the cluster shown in (B,B’), plotted as in Fig 3 (tracked SOP1, blue; neighboring cells, pink) with an additional plot corresponding to the maximum values measured over time in any of the untracked cells of the cluster (orange curve; see movie 4). In this cluster, these max values were associated with a neighboring SOP (SOP2; white dot in B,C). The sudden drop in the maximal intensity value at 6.5h resulted from the fact that SOP2 was no longer scored as a neighboring cell (B’). D-E’) Snaphots of a CadmKate (D,E) GFP-Sc (D’,E’) movie showing a pair of SOPs in direct apical contact around SB (D,D’) which moved away from each other prior to mitosis through cell-cell rearrangements (E,E’).

### Symmetry breaking at low Notch activity level

We next examined the dynamics of Notch activity during lateral inhibition. To do so, we used a destabilized GFP (deGFP) expressed downstream of a Notch-Responsive Element (NRE- deGFP) (He et al., 2019). Unexpectedly, a strong NRE-deGFP signal was detected in the posterior region of the ADHN at ∼14hr APF (Fig 7A, movie 3; note, however, that posterior-most cells did not express NRE-deGFP, suggesting that Notch provides a negative template for the posterior-most SOPs that emerged first). This signal decreased over time and evolved to produce a salt-and-pepper pattern, with presumptive SOPs emerging as non-fluorescent cells, which extended over the central region of the ADHN (Fig 7B,C). Averaging the NRE- deGFP signal over the posterior part of the ADHN showed that the NRE-deGFP signal decreased to reach a minimum at around ∼21 hr APF (∼7 hr after the start of the movie) before increasing again (Fig 7D). This decrease in fluorescence signal reflected that the degradation rate of deGFP was faster than its synthesis rate. Since the deGFP protein has a measured half-life of ∼2 hr (He et al., 2019), a 2-fold decrease in fluorescence over a 2h period may suggest that Notch signaling is off. Interestingly, the rate of decrease was maximal at ∼17.5h APF, indicating that Notch activity was minimal around the time of SOP emergence (Fig 7E). Next, to study Notch dynamics with single cell resolution, we used the same approach as the one used for GFP-Sc, focusing on the anterior neurogenic region of the ADHN where no initial NRE-deGFP expression was detected. All nuclei were segmented using a nuclear RFP and presumptive SOPs (identified as NRE-deGFP low cells; Fig 7C) were backtracked (n=21; 2 movies).

**Figure 7:**
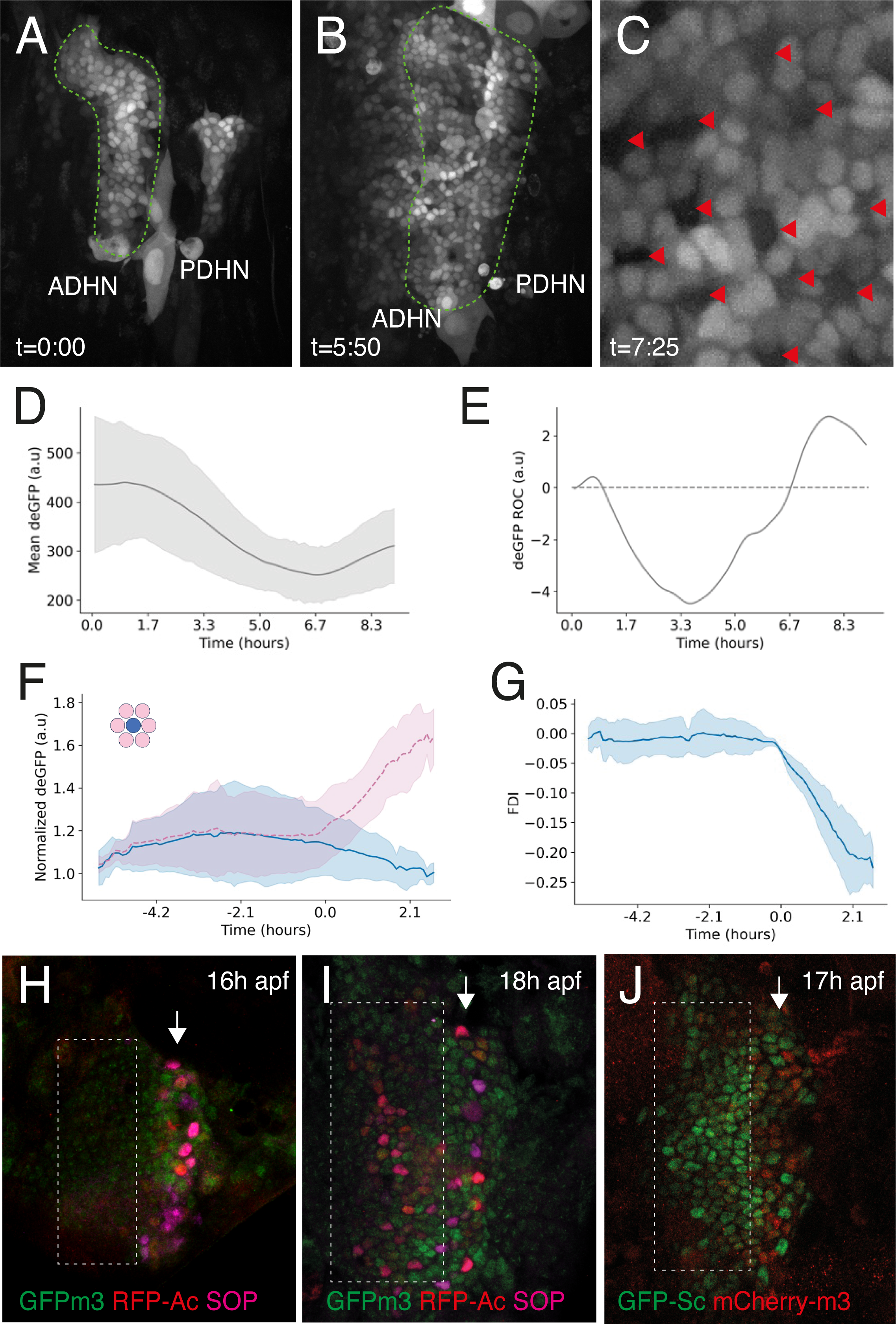
Dynamics of Notch activity. A-C) Notch signaling activity (NRE-deGFP, white) before and after SOP emergence (A,B; t=0 corresponds to ∼14h APF; see movie 5). Cells from the posterior region of the ADHN (green dotted line), as well as the PDHN, had received a Notch-activating signal prior to 14h APF (A). Following SOP emergence (B), Notch activity was detected in most cells of the central region of the ADHN (note that deGFP was no longer detected in the PDHN). SOPs were identified as NRE-deGFP low cells (C) and tracked using a nuclear nlsRFP marker (not shown). D) Temporal profile of the NRE-deGFP mean intensity signal from the posterior region of the ADHN, showing a gradual decrease until t=7h, corresponding to ∼21h APF. E) Temporal profile of the ROC of the NRE-deGFP signal. The maximal rate of decrease, corresponding to the minimum level of Notch activation, was observed at t=3.5h, corresponding to ∼17.5h APF. F) Normalized deGFP intensities over time for the 21 clusters registered in time using SB (t=0). The temporal profiles of NRE-deGFP in SOPs (blue) and non-SOP cells (pink) are shown. SB was detected at very low Notch activity levels. G) Temporal profile of FDI values for all registered clusters. An increase of fate divergence was observed from the time of SB onwards (FDI values were negative since deGFP intensity values increased in the SOP neighbors). H-J) The E(spl)-HLH factor m3 (GFPm3, green in H,I; mCherry-m3, red in J) was detected in the posterior region of the ADHN (white arrows) around SOPs (marked by Sens, magenta) at 16-17h APF. E(spl)m3 was detected in the central region (boxed area in I) soon after the onset of Sc (GFP-Sc, green in J) and Ac (RFP- Ac, red in H,I).

We next compared the relative levels of NRE-deGFP in each SOP relative to its closest neighbors, calculated a FDI and identified SB. We then registered all clusters relative to SB (defined as t=0) and plotted the temporal profile of NRE-deGFP expression (Fig 7F). We refer to this <u>SB</u> in <u>N</u>otch activity as SBN (to distinguish it from the SB determined using proneural dynamics). This showed that SBN occurred at very low Notch activity levels and that non-SOP cells showed increasing Notch signaling activity only after SBN (Fig 7G; note that FDI values were negative and decreasing as deGFP was expressed in non-SOP cells). Thus, this quantitative approach failed to detect a phase of reciprocal Notch signaling during which proneural cluster cells would both send and receive a Delta-Notch signal prior to SOP emergence. It also showed that SB was detected soon after the onset of Notch-mediated inhibitory signaling.

These conclusions were further supported by our analysis of the expression of the *E(spl)m3* Notch target gene. Indeed, a tagged version of endogenous *E(spl)m3*, GFPm3, was detected by antibody staining around emerging SOPs first in the posterior ADHN (16h APF; Fig 7H) then in the central ADHN domain (18h APF; Fig 7I; GFPm3 was not detectable in the abdomen of living pupae). Analysis of an mCherry-tagged E(spl)m3 showed that GFP-Sc was detected prior to E(spl)m3 in the central domain of the ADHN (Fig 7J; other E(spl)-HLH factors showed a similar spatial-temporal dynamics, see Fig S3). Thus, E(spl)m3 was detected after the onset of Sc expression and upon, but not prior to, SOP emergence, indicating that SOP selection in the abdominal epidermis took place at low signaling levels of Notch.

## Discussion

Quantitative live imaging of the proneural factor Sc allowed us to detect fate SB and monitor fate divergence during lateral inhibition in the pupal abdomen. This showed that SB takes place at low level of Sc. Following SB, the accumulation of Sc in presumptive SOPs was not associated with a concomitant decrease of Sc in all non-SOP cells. Instead, the mean level of Sc remained relatively constant whereas heterogeneity of Sc increased, indicating that signal-receiving cells were asynchronously excluded, as seen earlier in the pupal notum (Corson et al., 2017)(Couturier et al., 2019). We also found that cell-to-cell variations in Sc levels increased prior to SB. These variations positively correlated with fate divergence, and experimental increase of GFP-Sc heterogeneity resulted in a faster rate of fate divergence, confirming that initial differences in Sc levels become amplified via an intercellular feedback loop during lateral inhibition. The notion that cell-to-cell variability promotes fate divergence and increases patterning speed is not novel and applies to other juxtacrine signaling systems (Rudge and Burrage, 2008). Nevertheless, our observations are important as they provide experimental support for the intercellular feedback loop model (Collier et al., 1996; Heitzler and Simpson, 1991). Finally, SB was found to correlate in time with the SOP division, suggesting that a stereotyped program of gene expression and differentiation follows SB to direct asymmetric cell division (Schweisguth, 2015). Our analysis, however, failed to detect a phase of mutual inhibition before SB, during which proneural cluster cells are thought to express intermediate levels of Sc while experiencing low levels of Notch signaling (Cubas et al., 1991)(Campuzano and Modolell, 1992). While a phase of mutual inhibition is thought to be dispensable in contexts where binary fate decisions are strongly biased, as during neuroblast selection (Simpson, 1997), this phase is assumed to precede the selection of individual cells by Notch in contexts where stochastic and largely unbiased decisions produce salt-and-pepper patterns. Contrary to this view, we did not detect a phase of relatively uniform GFP-Sc expression prior to SB in the central region of the ADHN, and SB was instead detected at low Sc and Notch levels. Thus, lateral inhibition appeared to act early and fast, at low levels of proneural and Notch activities, in this tissue. Accordingly, initial bias in Sc expression might play a significant role in SB. The origin of the initial heterogeneity of Sc is unclear. In principle, cell-to-cell variations in Notch activity might provide a negative bias for the onset of Sc. Unfortunately, we could not test whether Notch activity anti-correlates with Sc levels prior to SB due to the low expression levels of Sc and Notch activity reporter at this stage. our observation that clusters producing late-emerging SOPs showed increased heterogeneity, presumably due to local Notch activation associated with the emergence of earlier SOPs in the vicinity, is consistent with this possibility. Other sources of heterogeneity may also be considered. For instance, in mouse intestinal organoids, cell-to-cell differences in mechanics have been involved in initiating a Delta/Notch signaling that drives SB (Serra et al., 2019). While a potential role of mechanics in regulating fate decision remains to be studied in the abdomen, our analysis of apical area indicated that the size of the cell-cell contact did not provide a clear bias for fate decision in this tissue. In other systems, non-genetic heterogeneities can result from cell-to-cell differences in the capacity of producing and maintaining a stable pool of active protein levels (Colman-Lerner et al., 2005). Future studies will address the basis of the early heterogeneity in Sc accumulation.

Our study also revealed that lateral inhibition did not faithfully single out SOPs in the pupal abdomen. Indeed, pairs of presumptive SOPs were unexpectedly found in direct cell-cell contact at the time of SB in ∼10% of the clusters. Thus, our observation suggested that the feedback loop may not be robust enough to faithfully impose an alternative fate choice and that SOP twins become rapidly deaf to inhibitory signals produced by their direct SOP neighbors. The basis for this patterning defect is not known. We speculate here that Notch cis-inhibition (del Álamo et al., 2011), which is known to down-regulate Notch and speed up fate divergence (Troost et al., 2023)(Sprinzak et al., 2011)(Sánchez-Iranzo et al., 2022), might facilitate the emergence of SOP twins. This hypothesis remains to be examined. Of note, pairs of emerging SOPs identified based on a proneural-responsive reporter had been reported earlier in the pupal notum of *Drosophila* (Cohen et al., 2010); however, this reporter was found later to be also expressed in a small number of non-SOP cells in this tissue (Corson et al., 2017), and it is therefore unclear whether pairs of SOPs are actually generated in the notum. Whatever the basis of this lateral inhibition defect in the abdomen, we observed that cell-cell rearrangements led to the separation of these SOP twins after SB, leading to pattern refinement. Since acquisition of the SOP fate is accompanied by changes in acto-myosin dynamics (Couturier et al., 2017) that might be associated with changes in adhesive properties. In this scenario, a fate-dependent change in acto-myosin dynamics would regulate differential cell adhesion to result in a loss of cell-cell contact between neighboring SOPs. Thus, changes in cell-cell adhesion downstream of Notch, as observed for instance in the pupal eye (Blackie et al., 2021), would contribute to patterning precision. Such a mechano-signaling feedback (Dullweber and Erzberger, 2023) might be one of the mechanisms needed to refine patterning.

## Acknowledgements

We thank Stein Aerts, Laure Bally-Cuif, Yohanns Bellaiche, Hugo Bellen, Francis Corson, Li He, Yang Hong, Romain Levayer, Juan Luna, Norbert Perrimon, Jean-Yves Tinevez, Alexis Villars, Flybase, Developmental Studies Hybridoma Bank and the Bloomington Drosophila Stock Center for reagents, resources, and advices. We thank the UtechS Photonic BioImaging (Imagopole; supported by France BioImaging (ANR-10–INBS–04) for use of a spinning-disk microscope. We thank R. Levayer for critical reading. Jang-mi Kim received a doctoral contract from Sorbonne University. Cara Picciotto received a LabEX REVIVE post-doctoral fellowship (ANR-10-LABX-0073). This work was funded by the Agence Nationale pour la Recherche (ANR-10-LABX-0073 and ANR-16-CE13-0003) and by the Fondation pour la Recherche Médicale (FRM-DEQ20180339219).

## Materials and Experimental Methods

### Flies

Sc was GFP-tagged at its N-terminus in the *scute^GFP^* knock-in line. This line was generated using CRISPR-mediated Homologous Recombination (HR) using *3xP3-RFP* as a selection marker (Corson et al., 2017). Since 3xP3-RFP gave a spotty signal in the abdominal nest, this marker, flanked by loxP sites, was removed using the Cre recombinase (BL-851). Proper excision was verified by gPCR. The *rfp^-^* version of *scute^GFP^* is fully functional and was used in this study. The RFP-Ac line carries a functional BAC-encoded version of the *ac* gene (Corson et al., 2017). The E(spl)m3-HLH factor was GFP-tagged at its N-terminus in the GFPm3 CRISPR knock-in line (Couturier et al., 2019), whereas the functional GFPm8, GFPmδ and GFPmγ factors were encoded in a transgenic BAC (Couturier et al., 2019). The Cherry-tagged version of E(spl)m3 was produced as a BAC transgene by recombineering in *E. coli* an *E(spl)-C* BAC (Chanet et al., 2009) as described for the GFPm3 BAC transgene (Venken et al., 2006)(Couturier et al., 2019) (cloning details available upon request). The mCherry-m3 BAC was integrated at attP site located in 99F8 (VK20 line). Injection was performed by BestGene Inc. (Chinmo, USA). The *sc[10-1] ac[3]* (BL-36541) mutant was used. The activity of Notch was monitored using the NRE-deGFP line (He et al., 2019). Different mRFPnls lines were used to mark nuclei. Apical junctions were tracked using versions of DE-Cad intracellularly tagged with GFP (Huang et al., 2009) or mKate (Pinheiro et al., 2017). SOPs were identified using a *neur-nlsGFP* transgene (Aerts et al., 2010). The silencing of GFP in marked flp-out clones was achieved using a flp-out strategy in pupae carrying hs-flp, nlsRFP and GFP-Sc on the X, P{y[+t7.7] v[+t1.8]=VALIUM20-EGFP.shRNA.3}attP40 (BL-41559) on the second chromosome and P{UAS-His3.3.mIFP-T2A-HO1}attP2 (BL-64184) and P{w[+mC]=AyGAL4}17b (BL-4413) on the third chromosome. Tubes with third instar larvae were heat-shocked (40min, 36.5°C) in a water-bath.

### Live imaging

Imaging of living pupae can be performed at high spatial resolution from 13h APF onwards, after the pupal case detaches from the pre-cuticle secreted by the developing imago, and after histolysis of abdominal muscles, preventing body movements and allowing for stable imaging. Pupae were staged at 0h APF and mounted for imaging by removing the pupal case using fine forceps at ∼14h APF. Prior to imaging, larvae and pupae were grown at 18, 21 or 25°C, but image acquisition was done at 23-25°C. Slight differences in the absolute timing of SB may have resulted from these varying conditions. Spinning disk microscopy was performed using either a Leica DMRXA microscope equipped with a 40x (PL APO, N.A. 1.32 DIC M27) objective, a Yokogawa CSU-X1 spinning disk, a sCMOS Photometrics PRIM95B camera, 491/561/642 lasers and the Metamorph software, or a Nikon Ti2E microscope equipped with a 40x (N.A. 1.15 Water WD 0.6) objective, a Yokogawa CSU-W1 spinning disk, a sCMOS Photometrics PRIM95B camera and 488/561/640 lasers. Typically, we imaged the GFP every 5 mins with 20% laser power for 300ms and RFP every 2.5 mins with 9% laser power for 200ms. A z-stack of of ∼40 mm was imaged with Δz=1.33mm and 0.7mm for GFP-Sc and Cad-GFP movies, respectively. Reproducible dynamics of Sc was obtained over 10-16 hours of imaging.

### Image analysis

#### Segmentation of nuclei

The nuclear channel of the GFP-Sc nlsRFP movies was preprocessed to correct the contrast and denoised using Noise2Void (Krull et al., 2019). The 3D segmentation of nuclear boundary was performed based on a custom-made program (Corson et al., 2017). Undesired structures (dead corpses, non-epithelial cells) among the segmented objects were filtered using different criteria (object volume, object intensity, object solidity). Nuclei belonging to the *gfp* RNAi clone were identified using intensity-based thresholding of the His3.3-mIFP signal.

#### Tracking and analysis of SOP neighbors

The data was fed to Mastodon (https://github.com/mastodon-sc) for semi-automatic tracking of SOPs from their birth to their division (when available). The presumptive SOPs were identified as high GFP (GFP-Sc movies) or low GFP cells (NRE-deGFP movies) on maximal z-intensity projection at late stages. The six closest neighbors of SOPs were automatically detected in 3D based on their physical distances to SOPs and errors (due to the presence of dead corpses, over-segmentations, non-epithelial cells) were manually corrected using a customized web application (https://gitlab.pasteur.fr/4dunit/tracked-nuclear-neighbor-analyzer). The GFP signal was normalized by dividing the value measured in a single nucleus to the mean value of the signal measured at each time point in the same channel in the entire image stack (given the low intensity of the GFP-Sc signal, this value should closely reflect the autofluorescence noise).

#### Kymograph

The GFP-Sc kymograph was computed by dividing the AP axis into continuous bins with width = 5 pixels. Then, the mean GFP-Sc of all segmented nuclei within each bin at a given time is calculated and normalized by the mean GFP-Sc of the image stack at that time.

#### Segmentation of apical junctions and tracking of SOPs

Images were processed using max z-intensity projection for the most apical 3-4 z stacks to limit the projection to Cadherin signal. Probability map of cells was generated using TrainableWeka (Arganda-Carreras et al., 2017), segmented automatically and manually corrected on Tissue Analyzer (Aigouy and Prud’homme, 2022). SOPs were semi-automatically tracked on Tissue Analyzer.

#### Data processing

The fate difference index (FDI) of a SOP at a given time *t* is defined as:

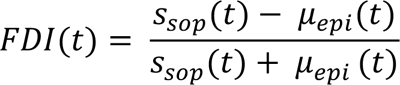

where *S_sop_*(*t*) is the GFP-Sc or NRE-deGFP of SOP and *μ*_*epi*_(*t*) is the mean GFP-Sc of its six nearest neighbors at the time *t*.

For the movies of RNAi cells clones, the SOP’s neighbors expressing RNAi are excluded when computing the FDI.

The point of symmetry breaking (SBP) of GFP-Sc is defined as the time *t*_!_ from which the FDI is greater than or equal a given threshold ε:

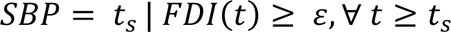

For the SBP of NRE-deGFP, it is defined as:

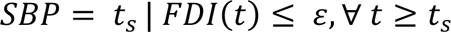

*ε* is set as 0.005 for GFP-Sc movies and as -0.025 for NRE-deGFP movies.

The heterogeneity is computed among cells in cluster (SOP and its six nearest neighbors). It is defined as:

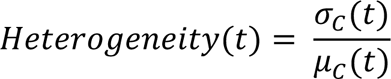

where σ_*c*_(*t*) and *μ_c_*(*t*) are the standard deviation and mean of GFP-Sc or NRE-deGFP of cells in the cluster at the time *t*. Heterogeneity at SB was calculated as the average over a 20min time interval centered at SB. The ROC of the FDI was calculated within a 20min time interval starting at SB.

Following segmentation of the apical Cad-GFP signal using Tissue Analyzer, apical area values and cell-cell junctions IDs were collected. The cells in direct contact with the tracked SOPs were identified using shared cell-cell junction ID at each time. This also provided the number of SOP neighbors (5.8 +/- 1.1, n=35). Apical area of tracked SOPs was then divided by the mean of apical area of neighbor cells. Dividing cells, that have large apical areas, were removed from analysis. To do so, cells with apical areas greater than 400 microns were excluded. Time registration of the different clusters was performed using the SOP division as a reference time point.

### Statistical analysis

Linear least-square regression was used to estimate the linear correlation between two datasets and the slope was tested by Wald test. Pearson correlation analysis was used to measure correlation strength (R). Mean and standard deviation were used in time-series plots.

### Imunostaining and microscopy

Dissection of staged pupae and antibody staining were performed using standard procedures. Briefly, nota were dissected from staged pupae using Vannas micro-scissors, fixed in paraformaldehyde (4% in PBS 1x) and incubated in PBS 1x with 0.1% Triton X100 and primary antibodies for 1.5 hr at room temperature. The following antibodies were used: goat anti-GFP (Abcam ab6673, 1:1000), rabbit anti-DsRed (Clonetech #632496, 1:500), guinea-pig anti-Sens (from H. Bellen, 1:2000) (Nolo et al., 2000), mouse anti-Cut (2B10, from DSHB, 1:100) and mouse anti-Hnt (1G9, from DSHB, 1:50). Secondary antibodies were from Jackson’s laboratories. Following washes in PBT, nota were mounted in 4% N-propyl-galate, 80% glycerol.

Images were acquired using a confocal Zeiss LSM780 microscope with 63x (PL APO, N.A. 1.4 DIC M27) and 40x (PL APO, N.A. 1.32 DIC M27) objectives.

Adult flies were imaged using a Zeiss Discovery V20 stereo-macroscope using a 1.0X (PlanApo S FWD 60mm) objective.

## Supplemental data

**Figure S1: role of Ac and Sc in SOP determination**

A) Dorsal view of an adult *sc^10-1^ ac^3^* fly showing a strong loss of sensory bristles in the abdomen, resulting from a loss of SOPs in pupae.

B) The loss of *sc* activity that resulted from the RNAi-mediated silencing of *scute^gfp^* in clones of cells did not prevent the formation of SOPs. Clones were identified by the loss of GFP-Sc (anti-GFP, green; nlsRFP, blue) in 22h APF pupae. Hindsight (Hnt) and Cut (both red) were used as SOP markers.

C,D) The silencing of *gfp* in clones in *sc^gfp^* pupae did not affect clone size (C) but strongly biased the SOP fate decision along clone borders (C; p=0.0003, unpaired t-test), as wild-type SOPs were more likely to adopt the SOP fate that cells with low sc levels (D).

**Figure S2: FDI threshold value**

A) Schematic representation of the approach. FDI values were measured on randomized clusters composed of non-SOP cells after the masking of tracked SOPs from segmented images.

B) Plot of the mean and sdt of FDI values measured over time in these randomized clusters. This analysis was performed in the central ADHN domain (n=3 movies).

**Figure S3: expression of E(spl)-HLH in the pupal abdomen**

The expression patterns of the E(spl)-HLH factors m8 (GFPm8, A), mγ (GFP mγ, B) and mδ (GFP mδ, C) were examined in the ADHN of fixed RFP-Ac (red) pupae. At 18h APF, these factors were detected in cells located close to emerging SOPs expressing Ac, notably along the posterior edge of the ADHN (white arrows), but were not detected before the local onset of Ac expression (see the central part of the ADHN, boxed in A-C).

**Movie 1: Live imaging of GFP-Sc in wild-type pupae**

Dynamics of GFP-Sc (green) in the pupal abdomen (nuclear RFP, red). This movie is a maximum projection view of a (x,y,z,c,t) movie. The ADHN and PDHN are seen as expanding islands of small proliferating cells surrounded by large larval epidermal cells that are progressively eliminated. A row of posterior SOPs appeared first before a salt-and-pepper pattern of SOPs emerge in the central region of the ADHN. Time is in h:min.

**Movie 2: Live imaging of GFP-Sc in mosaic pupae**

Dynamics of GFP-Sc (green) in the pupal abdomen of genetically mosaic pupae carrying clones of cells expressing a *gfp* shRNA construct and a Histone3.3-mIFP marker (blue; nuclear RFP, red). Time is in h:min.

**Movie 3: Apical view of the proliferating ADHN epithelium**

Live imaging of CadGFP allowed the back-tracking of SOPs, identified at mitosis using the pneur-nlsGFP signal (the strong SOP-specific nlsGFP signal filled the entire volume of dividing SOPs following nuclear membrane breakdown). Time is in h:min.

**Movie 4: An example of an emerging pair of SOPs**

Dynamics of GFP-Sc (green) and Cad-mKate (red) showing a pair of SOPs (indicated by yellow dots at the start/end of the movie) that remained in direct contact from t=0:00 to t=4:20. SB is predicted to take place around t=1:30 (both SOPs divide around ∼6h; not shown). Time is in h:min.

**Movie 5: Live imaging of a Notch activity reporter**

Live imaging of NRE-deGFP (green; nlsRFP, red). Cells in the posterior region of the ADHN (had received a Notch-activating signal prior to the onset of imaging. Notch activity became detectable in the anterior cells of the central region of the ADHN as SOPs (identified as NRE- deGFP low cells) emerged. Note that a small number of larval ectodermal cells were also positive for NRE-deGFP. Time is in h:min.

