## Supplementary figures and images for "Symmetry breaking and fate divergence during lateral inhibition in *Drosophila*"

### Figure S1

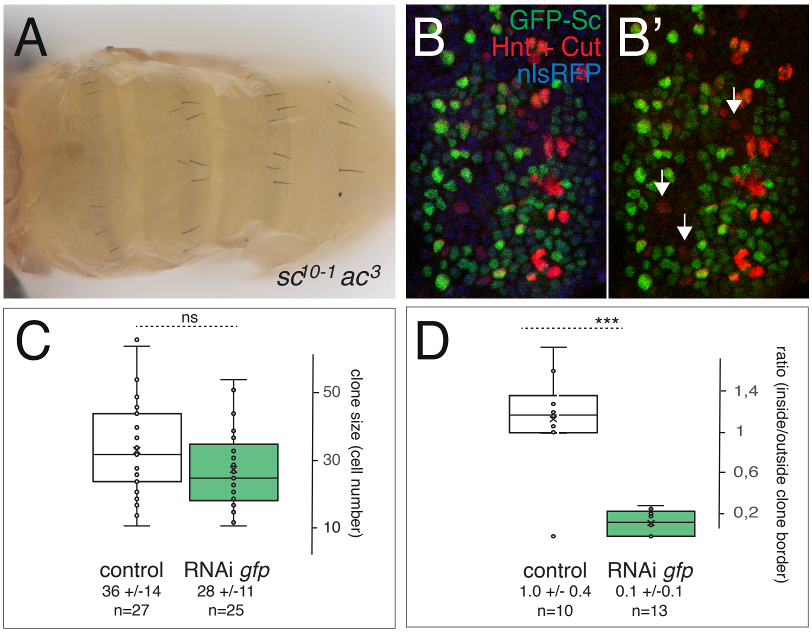

### Figure S2

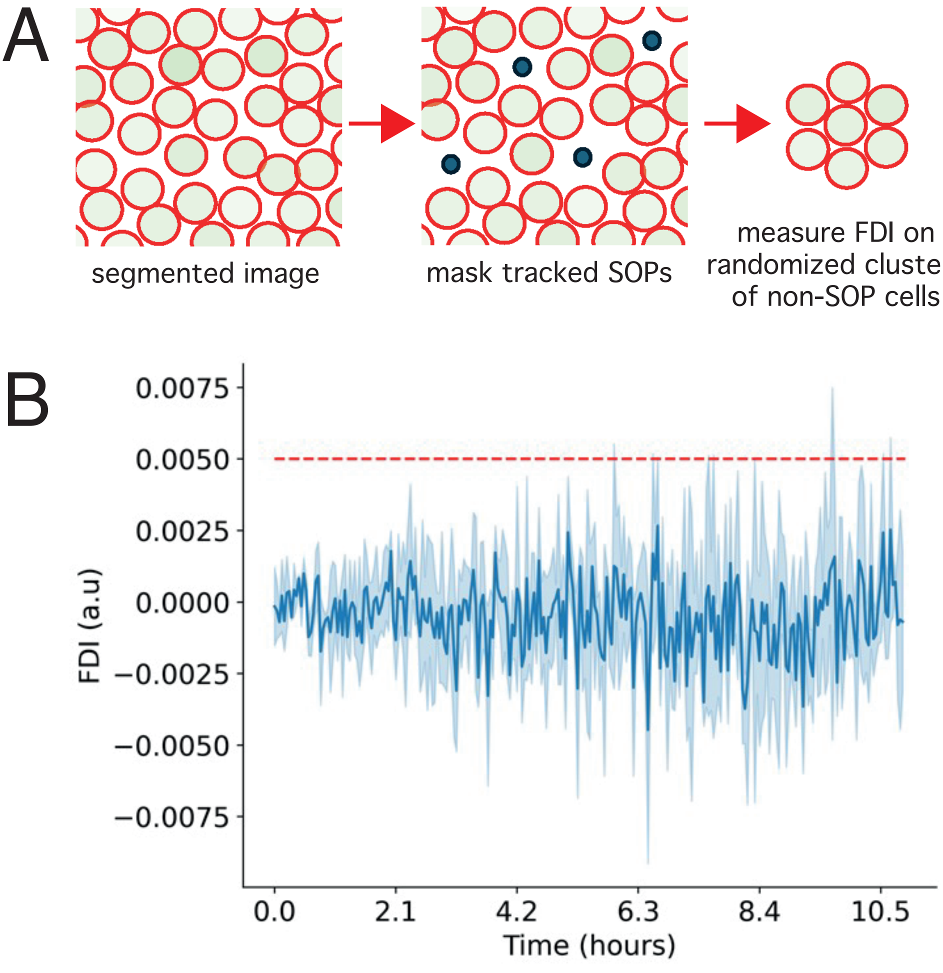

### Figure S3

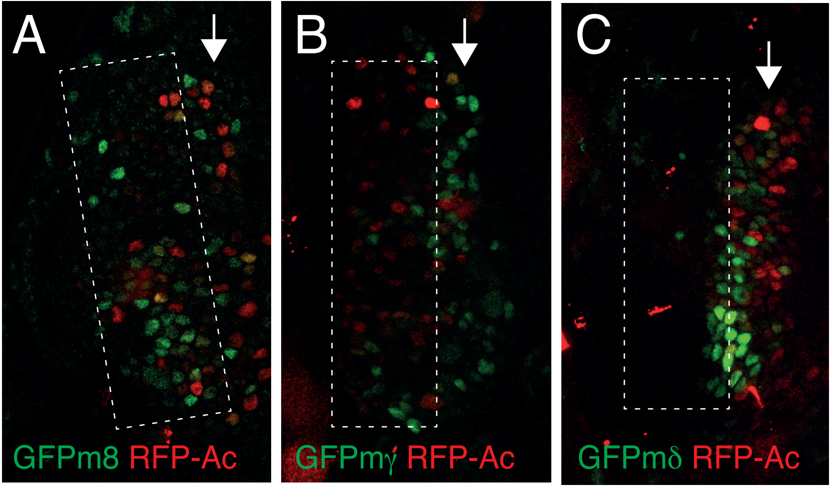
